# Inference of Causal Interaction Networks of Gut Microbiota Using Transfer Entropy

**DOI:** 10.1101/2024.04.15.589473

**Authors:** Chanho Park, Junil Kim, Julian Lee

## Abstract

Understanding the complex dynamics of gut microbiota interactions is essential for unraveling their influence on human health. In this study, we employed transfer entropy analysis to construct a causal interaction network among gut microbiota genera from the time-series data of bacterial abundances. Based on the longitudinal microbiome data from two subjects, we found that the constructed gut microbiota regulatory networks exhibited power-law degree distribution, intermediate modularity, and enrichment of feedback loops. Interestingly, the networks of the two subjects displayed differential enrichment of feedback loops, which may be associated with the differential recovery dynamics of the two subjects. In summary, the transfer entropy-based network construction provides us with valuable insights into the ecosystem of gut microbiota and allows us to identify key microbial hubs that play pivotal roles in shaping the microbial balances.

## Introduction

The human microbiota, particularly the gut microbiota, plays a crucial role in maintaining our health. This vast and dynamic assembly of microorganisms inhabits various parts of our body, with the gut microbiota being one of the most influential due to its extensive interactions with human physiology. Many studies has shed light on the profound impact the gut microbiota on human health [1–9]. They assist in the digestion and fermentation of the food, enabling the extraction and synthesis of vital nutrients and energy from the food we consume [5,6]. Beyond their role in nutrition, these microbial communities serve to protect against pathogens [7,8], regulate immune function, and strengthen the biochemical barriers of the gut and intestine [9].

Various bacterial species forming the microbiota interact with each other for their survival. In particular, the network of causal relationships among microbiota is of significant importance as it provides insights into the complex and dynamic ecosystem of microorganisms that are critically important in the function of any biological community. These networks can help us understand how commensal and pathogenic microbiota modulate host signaling and have broad cross-species consequences.

Early studies on microbiota interactions relied on analyzing correlations between them, which produce only undirected interaction networks [10–16]. More detailed information on the nature of interactions can only be obtained by analyzing the time series of bacterial abundances, where directed graphs are obtained [17–20]. While significant progress has been made, the rigorous inference of causal relationships between bacterial species using information theory remains an underexplored area. In this study, we address this gap by constructing a causal interaction network employing transfer entropy (TE) [21], an information-theoretic measure for determining causality between variables.

TE has been used for inferring causal relationships within dynamic systems, including the functional connectivity of neurons [22–24], social system [25], and transcriptional regulation [26–28], by measuring the information transferred from one agent in the interacting network to another. In particular, we used the TENET (Transfer Entropy-based causal gene NETwork) algorithm [26], which is designed to construct gene regulatory networks (GRNs) from single-cell RNA sequencing data. TENET has demonstrated superior performance in identifying key regulatory transcription factors. Employing TENET, we built the causal interaction network of gut microbiota, by analyzing one-year time series of microbiota data from two individuals [29]. The resulting microbiota causal network revealed: 1) a power-law degree distribution with a few super-hub nodes; 2) intermediate modular structure characterized by phylum; and 3) an enrichment of feedback loops. Key bacterial genera, such as *Acrobacter* and *Clostridium*, were pinpointed as central hubs or connectors in these networks, with associations to the hosts’ health outcomes or environmental interactions. These findings suggest that TE is a valuable tool for identifying critical regulatory genera within gut microbiota, which could be potential candidates of therapeutic targets.

## Materials and Methods

### Processing of time-series data

One year time-series of gut microbiota for two people have been obtained in David *et al*. [29]. David *et al*. generated a sequencing data of V4 region of the 16S ribosomal RNA gene subunit on DNA obtained from stool samples collected daily from two subjects, subject A for 341 days and subject B for 192 days. The raw data resulting from this work are available in the European Nucleotide Archive (ENA) (https://www.ebi.ac.uk/ena/browser/view/PRJEB6518) maintained by the European Bioinformatics Institute (EBI). Post-processed time-series data are also available at MGnify database (https://www.ebi.ac.uk/metagenomics/studies/MGYS00001278#analysis), at the level of 32 phyla and 879 operational taxonomy units (OTUs). We used the time-series data for the OTUs and grouped them according to genera, resulting in the time-series abundances of 667 genera (Supplementary Table S1 and S2).

### Construction of gut microbiota regulatory network

TENET algorithm [26] computes transfer entropy for a given pair of components in an interaction network, when time-series for these components are given. The transfer entropy is defined as [21]

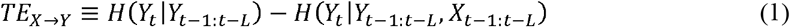

where *X*_*t*−1:*t*−*L*_ and *Y*_*t*−1:*t*−*L*_ denote the time series for the time-points *t*-*L, t*-*L*+1, … *t*-1, and *H*(*X*) is the Shannon’s entropy:

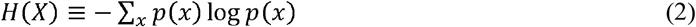

with *p*(*x*) being the probability that *X* takes the value *x*. The transfer entropy *TE*_*X*⟶*Y*_ measures the uncertainty on *Y* removed by gaining the information on the history of *X*, and estimate the information flow from *X* to *Y*. For estimating conditional entropy, TENET utilized a kernel density estimator, a non-parametric approach to estimate the probability density function of a random variable [26]. For simplicity, we will consider only the effect of the immediate past of the current time-point and use *L* = 1. We chose *L* = 1 since TENET results was almost similar when *L* > 1 is applied in the original TENET study on single cell RNAseq [26].

TENET [26] was downloaded from github.com/neocaleb/TENET and applied to the time-series microbiota profiles at the genus level for each subject individually. We then represented the microbiota regulatory network as a directed binary graph, where a node represents a genus, and there is a directed edge from a node A to a node B if *TE*_*A*⟶*B*_ is significantly large. In order to assess the significance of a TE value, we assumed that the values of TE are distributed according to a normal distribution and performed a one-sided z-test. The *TE*_*A*⟶*B*_ from A to B was considered significantly large only if its false discovery rate (FDR) [30] obtained from the z-test was below the threshold θ. Even if *TE*_*A*⟶*B*_ is significantly large, we did not draw a directed edge from A to B if this is due to an indirect effect, where there is a node C such that A affects C and C and B, but there is no direct causal effect of A on B. We assume such an indirect influence of A on B when we found C such that *TE*_*A*⟶*B*_ is less than the minimum value of *TE*_*A*⟶*C*_ and *TE*_*C*⟶*B*_. In this case, we did not draw a directed edge from A to B in the final graph. We repeated this trimming process until no further eliminations is necessary.

To test the robustness of the properties of the directed binary network with respect to the FDR threshold θ, we used both θ=0.01 and θ=0.05. The constructed networks were visualized using Cytoscape 3.9.1 with “Edge-weighted Force directed” and “Group Attributes” layouts [31].

### Power-law degree distribution

Many biological networks follow power-law degree distribution [32]. We constructed the degree distribution of each network and fitted the distribution to the power-law equation:

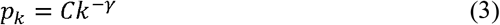

where *p*_*k*_ is the distribution or probability of degree *k, C* is a normalization constant ensuring that the sum of all probabilities *p*_*k*_ over all *k* equal *1*, and *γ* is the degree exponent. A network is called scale-free when it has a power-law degree distribution with 2<*γ*<3. The power-law degree distributions and degree exponents were obtained using the python package NetworkX.

### Leiden clustering

We used Leiden algorithm for community detection of network [33]. Although Leiden clustering algorithm is mainly designed for undirected graphs, we chose Leiden algorithm instead of algorithms for directed graphs, such as directed Louvain algorithm, since Leiden algorithm is more improved algorithm in detecting well-connected communities [33]. We also performed directed clustering Louvain to check that the results are similar to those obtained from the Leiden clustering. A general purpose of community detection algorithms is identifying communities within a network that are more densely connected internally than with the rest of the network. These algorithms aim to maximize modularity M:

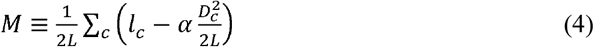

where *L* is the total number of links in the network, *l*_*c*_ is the number of links in community *c, α* is a resolution parameter, and *D*_*c*_ is the sum of the degrees of the nodes in community *c*. We used *α* = 0.5.

The Leiden algorithm begins with each node in its own community and iteratively moves to improve the community structure. After the initial community detection, the algorithm refines the partition to ensure that nodes within each community are well-connected. Based on the refined partition, the algorithm aggregates the network, where each community is represented as a node. Then the algorithm proceeds iteratively, refining and aggregating until the quality of the community detection cannot be improved any further. Leiden clustering was performed using clusterMaker2 [34] application embedded in Cytoscape with the following options: resolution = 1, beta value = 0.01, number of iterations = 2. The chosen resolution parameter influences the size and number of detected communities. The beta value adjusts the algorithm’s sensitivity to network density, and the specified number of iterations limits the algorithm’s refinement process, ensuring computational efficiency while seeking optimal community detection.

### Hypergeometric test

After clustering the nodes in the interaction network into several communities (see Results), we investigated whether certain phylum is enriched in a given cluster. Let us denote the total number of nodes as *N* and the number of nodes belonging to a given phylum, say phylum *A*, as *M*, and also denote the number of nodes in a cluster, say cluster *i*, as *n*. If we randomly select *n* nodes from *N* nodes, the number of nodes that belong to the phylum *A*, denoted as *K*, is a random variable. The probability distribution *P*_*K*_(*k*) for *K* is given by the hypergeometric distribution:

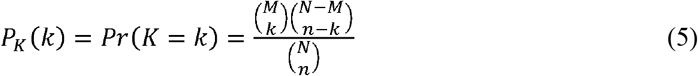

For the observed number 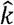 of nodes belonging to the phylum *A* in the cluster *i*, we also compute the *P*-value

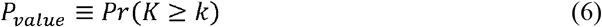

If the *P*-value is extremely small, we conclude there are significantly large number of nodes belonging to the phylum *A* than expected, and hence phylum *A* is enriched in the cluster *i*.

### Random rewiring of regulatory network

To measure the statistical significancy of the strength of regulations between phyla, we performed degree-preserving random rewiring using “rewire” function implemented in a R package “igraph”. We generated empirical distributions of the number of links between phyla by randomly rewired regulatory networks 1,000 times. For the observed number 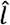 of links from phylum *A* to *B*, we calculate the *z*-score

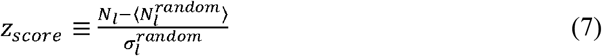

where *N*_*l*_ is the number of links from phylum A to *B*, and 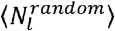 and 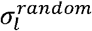 are the mean and standard deviation of links in the randomly rewired networks.

### Centrality measures

The out-degree and in-degree of a node *i* is defined as the number of out-going edges and the number of in-coming edges of the node *i*, respectively.

The betweenness centrality of a node *i* is defined as

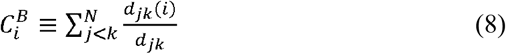

where *N, d*_*jk*_, and *d*_*jk*_ (*i*) denote the number of nodes, the number of shortest paths between node *j* and *k* and the number of shortest paths between node *j* and *k* which include node *i* in the path, respectively.

The closeness centrality of a node *i* is defined as

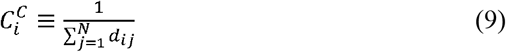

where *N* and *d*_*ij*_ denote the number of nodes and distance between node *i* and *j*, respectively.

### Network motifs

Network motifs are defined as patterns of interconnections observed more frequently than in an ensemble of randomized networks [35]. We searched for patterns involving two-node and three-node motifs using NetMatch [36] a Cytoscape plugin application.

The statistical significance of each motif was determined through z-scores, calculated by comparing the observed frequency against a null model. This model was generated by randomly shuffling the network’s edges 10,000 times, ensuring the degree distribution was maintained. To compare the z-scores of each motif between subject A and subject B, we calculated the z-scores 100 times.

## RESULTS

### Microbiota regulatory networks constructed by TENET display power-law degree distribution

With the time-course data on gut microbiota abundance, we calculated TE values for every pair of genera (Supplementary Tables S3-S4). Based on the TE matrix, we constructed gut microbiota regulatory networks for two subjects using TENET with θ = 0.01 (Supplementary Tables S5-S6). After trimming indirect edges, we measured the network density ρ ≡ *N*_*e*_ / *N*_*n*_ (*N*_*n*_ − 1), where *N*_*e*_ and *N*_*n*_ are the number of directed edges and the nodes, respectively. Since the network density ρ is the ratio of the actual number of directed edges to the maximum number of possible directed edges, 0 ≤ ρ ≤ 1. We found ρ = 0.012 for subject A and ρ = 0.021 for subject B, indicating sparsely connected structures. The degree distribution of both networks followed a power-law with degree exponents of *γ* = 0.913 for subject A and *γ* = 1.115 for subject B (Figures 1A and 1B), representing the existence of a few super-hub nodes connected with almost every other node, and a large number of nodes with a small number of edges. This sparsity and power-law degree distribution property persisted in the networks constructed with θ = 0.05 (Supplementary Table S7-S8 and Figure 1C and 1D). The goodness of fit of the power-law distribution was confirmed by comparing with exponential and log-normal positive distributions (Supplementary Table S9).

**Figure 1.**
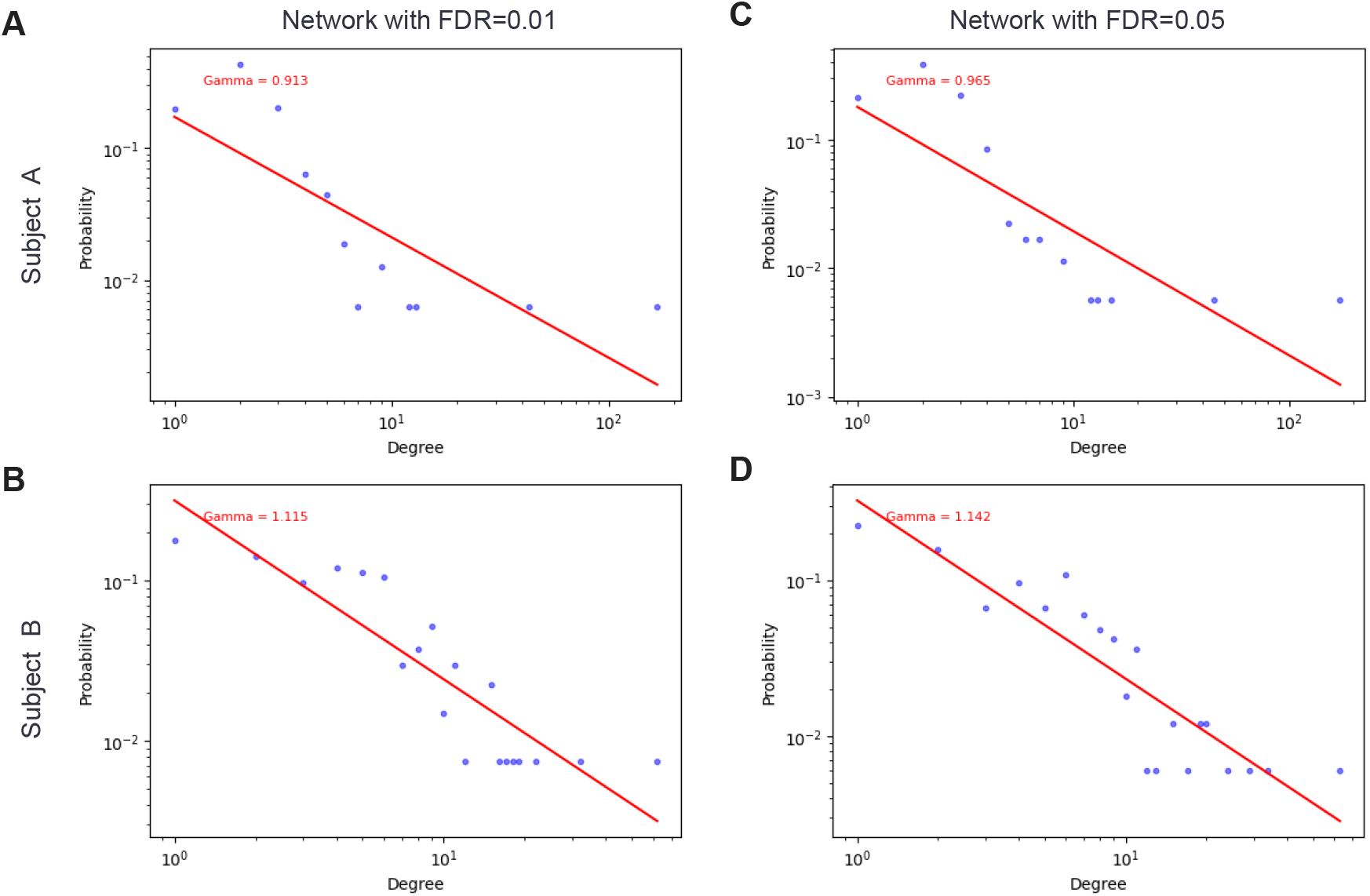
Gut microbiota regulatory networks display power-law degree distribution. (**A-B**) Degree distribution of gut microbiota regulatory networks for subject A (**A**) and B (**B**) with FDR threshold of 0.01. (**C-D**) Degree distribution of gut microbiota regulatory networks for subject A (**A**) and B (**B**) with FDR threshold of 0.05.

We then applied Leiden clustering algorithm to discern if these networks exhibit a modular organization [33]. The visualization of the clustering outcomes for subjects A and B is provided in Figures 2A and 2B, respectively. The modularity scores M were 0.288 for subject A and 0.35 for subject B, indicating intermediate modularity for both networks [37]. This modularity was also found in the constructed network with θ = 0.05 (M=0.317 for subject A and M=0.276 for subject B).

**Figure 2.**
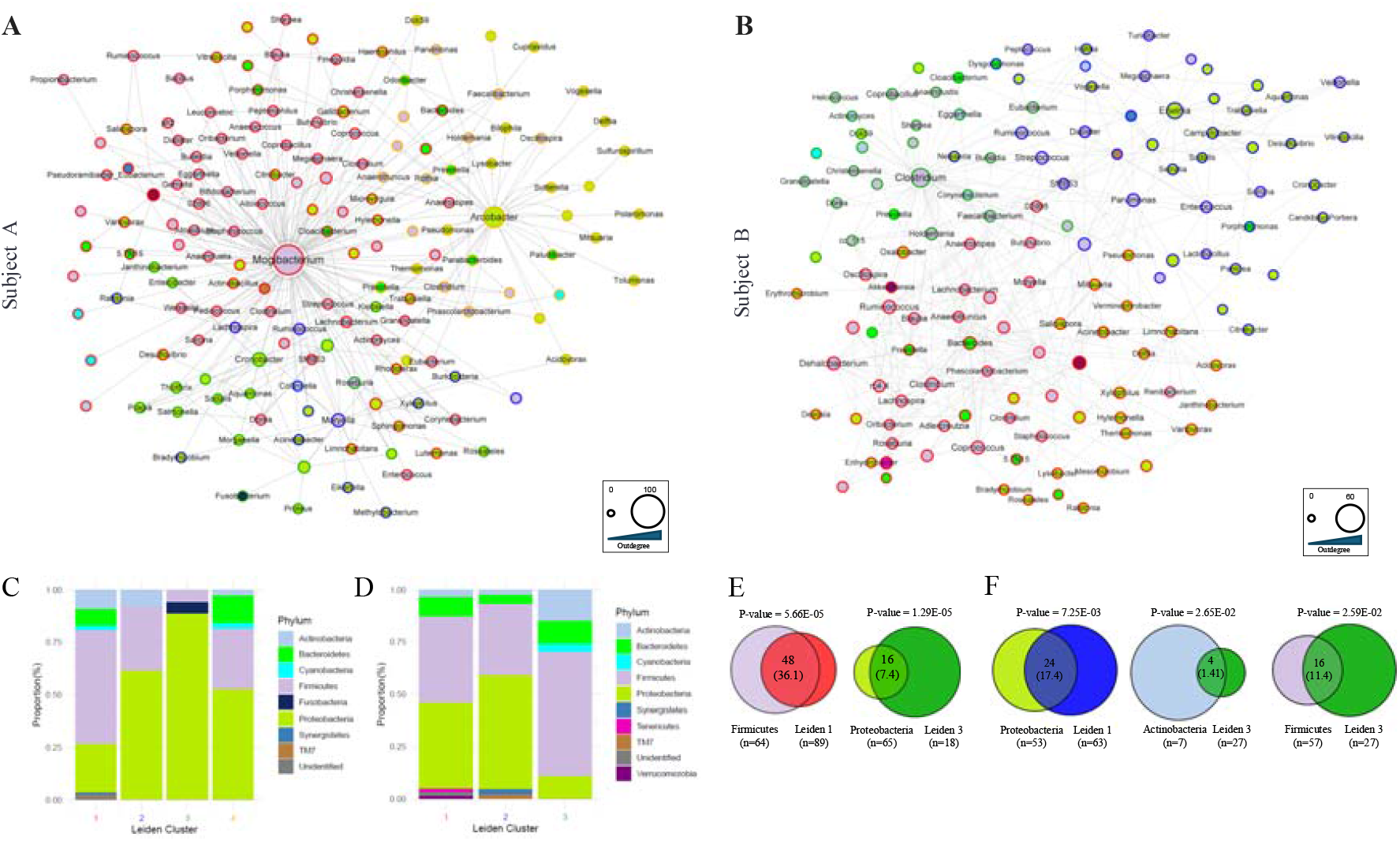
Gut microbiota regulatory networks exhibit modular structure. (**A-B**) Network representation of gut microbiota regulatory networks for subject A (**A**) and B (**B**) with “Edge-weighted Force directed” layout. The node fill color and border color represent phylum and Leiden cluster, respectively. The node size represents the out-degree. (**C-D**) The fractions of nodes belonging to distinct phyla for each Leiden cluster for subject A (**C**) and B (**D**). (**E-F**) The results the hypergeometric tests for the phylum compositions for several Leiden clusters in subject A (**E**) and B (**F**). The numbers in the intersection region denote the observed (upper) and expected (lower, presented in parenthesis) number of nodes.

Leiden clustering produced four and three clusters for the subject A and B, respectively. These are differentiated by the border colors of the nodes in Figures 2A and 2B. To associate each cluster with specific microbial phyla, we color-coded the interior of each node according to the phylum of the represented genus. We enumerated genera within each phylum for every cluster, with the compositional outcomes depicted in Figures 2C and 2D for subjects A and B, correspondingly. The color schemes for these figures are consistent with those in Figures 2A and 2B. Across all clusters for both subjects, two phyla - *Firmicutes* and *Proteobacteria* - were consistently present. Nevertheless, the exact phylum compositions differed by cluster. Hypergeometric testing, results of which are shown in Figures 2E, 2F, and Supplementary Table S10, identified phyla significantly overrepresented in certain clusters. For instance, *Firmicutes* were notably prevalent in Leiden cluster 1 of subject A (*p*=5.66E-5) and Leiden cluster 3 of subject B (*p*=2.59E-2), *Proteobacteria* were enriched in Leiden cluster 3 of subject A (*p*=1.29E-5) and Leiden cluster 1 of subject B (*p*=7.25E-3), and *Actinobacteria* were enriched in Leiden cluster 3 of subject B (*p*=2.65E-2). Additionally, some phyla appeared exclusively within specific clusters. For example, *Fusobacteria* were only found in Leiden cluster 3 of Subject A, *Synergistetes* were only found in Leiden cluster 1 of subject A and Leiden cluster 2 of subject B, and *TM7* were only found in Leiden cluster 1 of subject A and Leiden cluster 2 of subject B. Overall, these findings from our constructed networks reveal discernible communities characterized by unique phyla. The results from the Louvain clustering were similar (Supplementary Table S11).

### Random rewiring of the network revealed significant regulatory strengths between phyla

To assess the strength of regulation between phyla, we changed the layout of the networks using Cytoscape by grouping genera with same phylum (Figures 3A and 3B). Random rewiring of the networks was performed 1,000 times while preserving the degree distribution, providing z-values of the regulations between phyla (Figures 3C and 3D). In both subjects, regulations within *Firmicutes* and *Proteobacteria* were significantly enriched, while regulations between these two phyla were significantly depleted. This suggests that regulations within these phyla are more enriched compared to regulations between phyla. Interestingly, the regulations from *Fusobacteria* to *Proteobacteria* were significantly enriched in the network of subject A (Figure 3C), whereas the regulations within *Actinobacteria* and the regulations from *Tenericutes* to *Bacteriodetes* were significantly enriched in the network of subject B (Figure 3D). These results were consistently found in the constructed network with θ = 0.05 (Supplementary Figure 1). This suggests that *Fusobacteria, Actinobacteria*, and *Tenericutes* may play a differential role on the regulation of gut microbiota between the two subjects.

**Figure 3.**
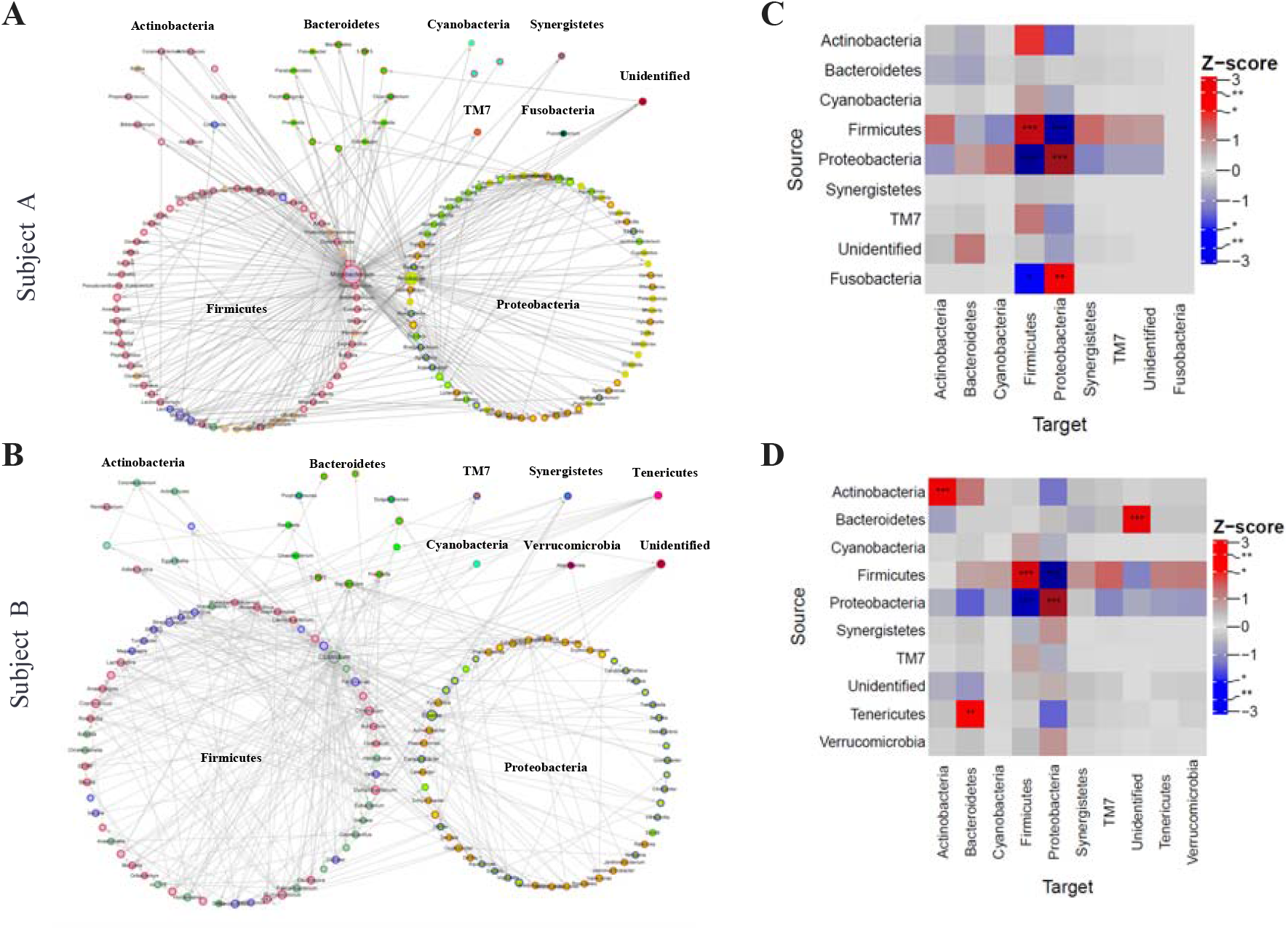
Random rewiring analysis of gut microbiota regulatory networks reveals significant regulatory relationships between phyla. (**A-B**) Network representation of gut microbiota regulatory networks for subject A (**A**) and B (**B**) with layout of grouping by phyla. The node fill color and border color represent phylum and Leiden cluster, respectively. The node size represents the out-degree. (**C-D**) Heatmaps of the z-scores of the number of regulations between phyla for subject A (**C**) and B (**D**). The networks were obtained using FDR threshold of 0.01.

### TENET identified key regulatory genera associated with personal experiences

In many network biology studies, there is a common hypothesis that certain key players, or ‘hub nodes’ play a critical role in maintaining the connections and flow within these networks [32,38,39]. In some studies, other measures of the importance of these hubs, such as their role as connectors or how centrally located they are in the network (i.e. betweenness or closeness), might be better at pinpointing critical proteins or genes [40–42]. To investigate this idea in the microbiota regulatory networks, we calculated four centrality measures: out-degree, in-degree, betweenness, and closeness (Figure 4). The centrality analysis revealed subject-specific key regulatory genera. In subject A, the genera *Mogibacterium*, belonging to the family *Mogibacteriaceae*, and *Arcobacter*, belonging to the family *Campylobacteraceae*, were consistently identified as key players across all four measurement methods (Figure 4A). However, in subject B, while two genera, *Clostridium*, belonging to the family *Ruminococcaceae*, was top first central in three of the methods, out-degree, in-degree, and betweenness centrality, it was not top first in closeness centrality (Figure 4B). Instead, two different genera *Dysgonomonas*, belonging to the family *Porphyromonadaceae*, and *Peptococcus*, belonging to the family *Peptococcaceae*, were top ranked in closeness. This indicates that *Dysgonomonas* and *Peptococcus* play a bridge role between modules instead of being a local hub. These key regulatory genera were also found in the regulatory networks constructed with an FDR threshold of 0.05 (Supplementary Figure 2).

**Figure 4.**
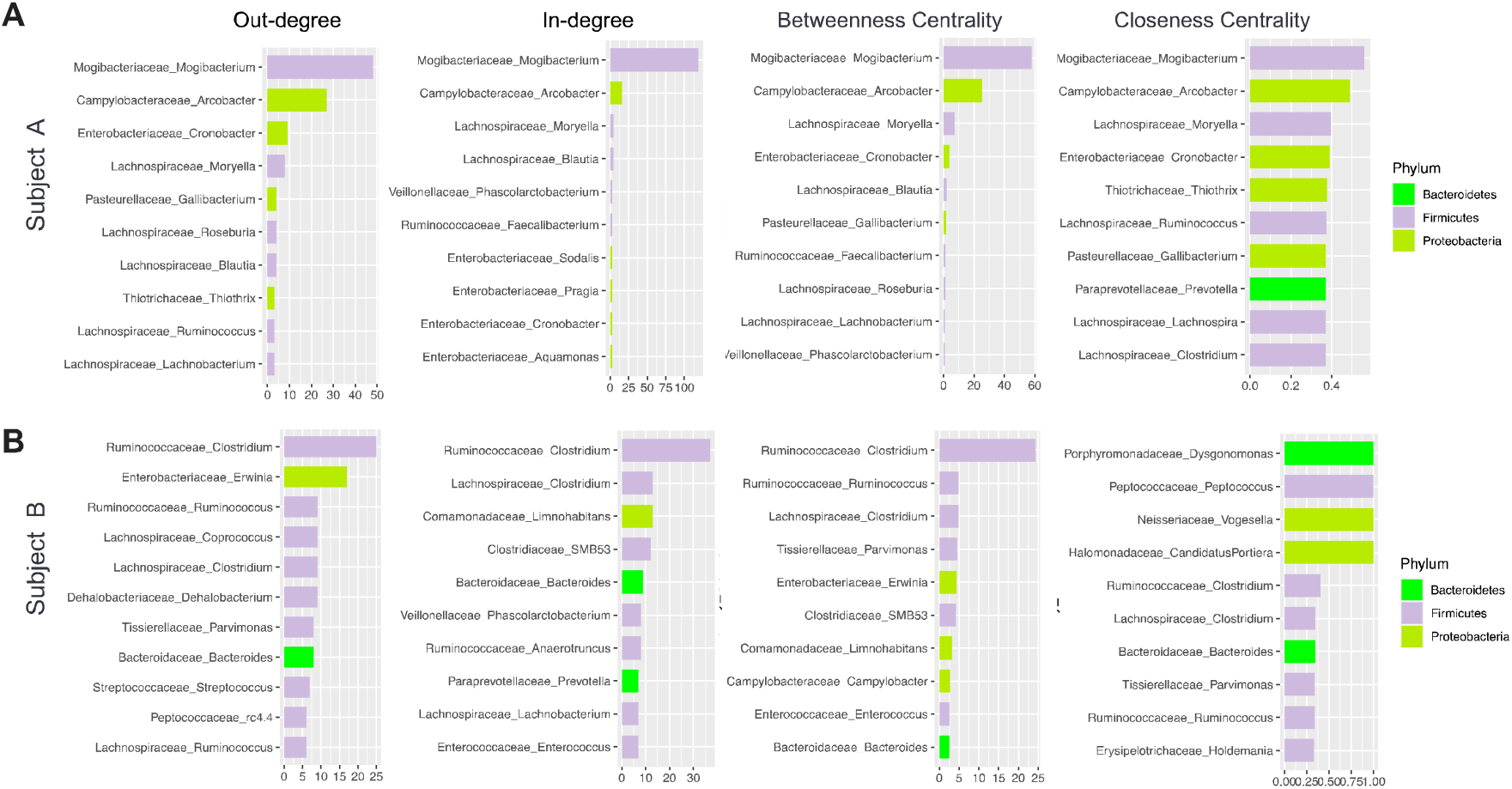
Centrality analysis of gut microbiota regulatory networks reveals candidates of key regulatory genera for regulating gut microbiota. (**A-B**) Four centrality measures (outdegree, indegree, betweenness centrality, and closeness centrality) of subject A (**A**) and B (**B**). The color for bar represents phylum. The networks were obtained using FDR threshold of 0.01. **course abundance of key regulatory microbiota**. (**A-B**) Time course abundance of *Acrobacter* in subject A (**A**) and *Clostridium* in subject B (**B**).

For the comparison, we explored two additional network inference algorithm, correlation network and SpiecEasi [43]. We calculated degree of the result networks since both algorithms provide undirected networks. For Subject A, the top 10 genus for the highest degree did not include microbe directly related to travelers’ diarrhea. In the case of Subject B from the correlation network, *Edwardsiella*, particularly *Edwardsiella tarda*, was noted for its relevance to food and waterborne infections. *E*.*tarda* is recognized for causing gastroenteritis and bacteremia, conditions closely tied to exposure to freshwater environments and seafood consumption [44] (Supplementary Figure 3).

The centrality analysis results lead us to propose that these central regulatory genera may influence the dynamic abundance of other genera, which can be linked to the individual characteristics or conditions of each subject being studied. Subject A experienced traveler’s diarrhea (TD) while traveling to Southeast Asia. *Acrobacter*, one of the key regulatory genera displayed peak abundance right after the traveling period (Figure 5A), representing regulatory role of the genus. *Acrobacter* is a microbial genus belonging to the *Proteobacteria* phylum and is associated with TD, which is frequently encountered in Southeast Asia [45–47]. Specifically, *A. butzleri*, one of the *Acrobacter* species has been extensively studied and demonstrated to have considerable importance for TD [48– 51].

**Figure.**
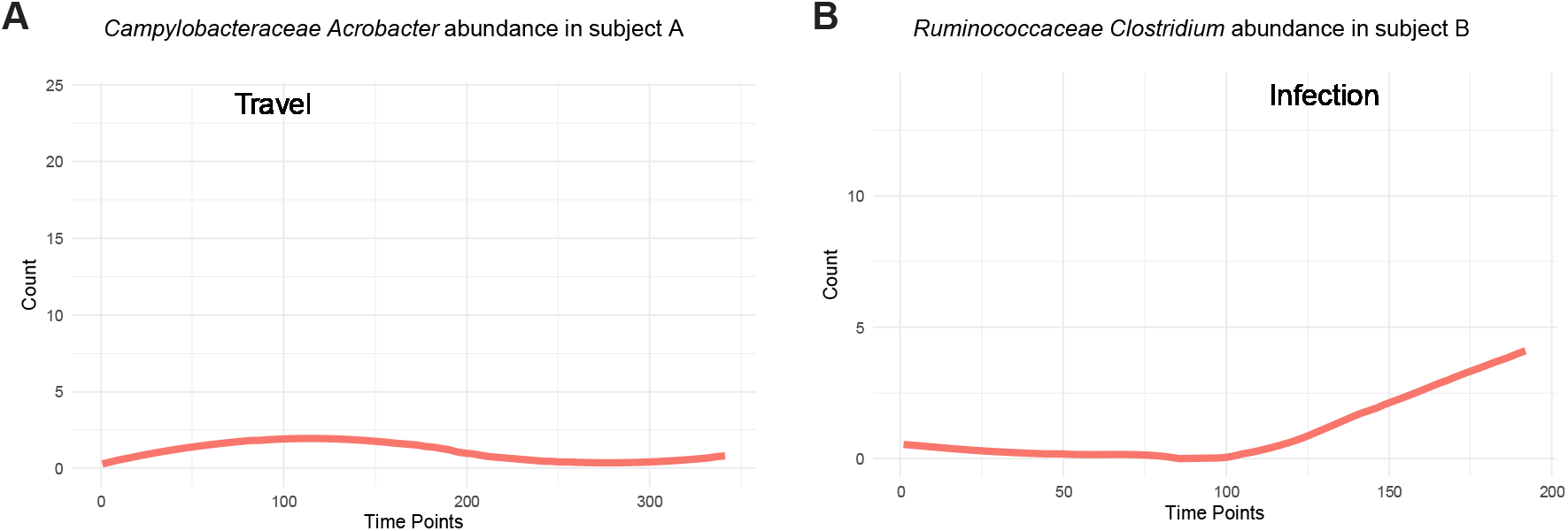
Timecourse abundance of key regulatory microbiota. **(A-B)** Time course abundance of *Acrobacter* in subject A **(A)** and *Clostridium* in subject B **(B)**.

Subject B experienced food poisoning, which is consistent with an increase in sequencing reads of *Enterobacteriaceae Salmonella*, during the infection period. However, the network for subject B indicates that *Clostridium*, part of the *Firmicutes* phylum, displayed the highest out-degree and in-degree, the second highest betweenness centrality, and the fifth highest closeness centrality. Notably, *Salmonella* did not appear in the list of top 10 key genera based on these centralities. In the original longitudinal microbiome study, the authors noted a significant increase in an OTU cluster, predominantly composed of the *Firmicutes* phylum including *Clostridium* following the infection.

This cluster maintained its elevated abundance, implying a loss of compositional stability in subject B [29]. We corroborated this significant increase in the abundance of *Clostridium* after infection (Figure 5B). Our finding of *Clostridium* as a key regulatory genus represents that this genus may influence the abundances of other species to counteract the external infection by *Salmonella*.

### Differential enrichment of feedback loops in two subject’s networks

In the original longitudinal microbiome study, the authors proposed that the microbiota of the two subjects demonstrated different recovery dynamics [29]. The microbiota community from subject A showed reversible dynamics after traveling abroad, whereas that from subject B showed irreversible changes after an enteric infection. To further investigate the differential recovery dynamics, we searched for network motifs, including a two-node feedback loop and thirteen possible three-node motifs, based on the relevance between feedback loops and steady-state dynamics [52,53]. The results showed differential enrichment of feedback loops between the two subjects (Figure 6). Both the two-node feedback loop and three three-node motifs associated with the two-node feedback loops (M11, M12, and M13) were significantly enriched in both networks. However, the significance levels of these four motifs were much higher in subject B’s network (z-score and p-value = 8.03 and 4.44E-16 for two-node feedback loop, 8.28 and 1.11E-16 for M11, 6.3 and 1.49E-10 for M12, and 11.1 and 6.27E-29 for M13) than in subject A’s network (z-score and p-value = 4.45 and 4.29E-6 for two-node feedback loop, 3.63 and 1.42E-4 for M11, 3.56 and 1.85E-4 for M12, and 3.4 and 3.37E-4 for M13). On the other hand, motif M9 representing cascade was only observed in Subject B’s network, with a z-score of −8.43 and a p-value of 1.48E-16. These differential enrichment and depletion of motifs between two subjects’ networks, was consistent when we repeated motif analysis 100 times (Supplementary Figure 4). This finding suggests that feedback loops may be a critical factor in the differential recovery dynamics. Consistent results also obtained from the constructed networks with an FDR threshold of 0.05 (Figure 6).

**Figure 6.**
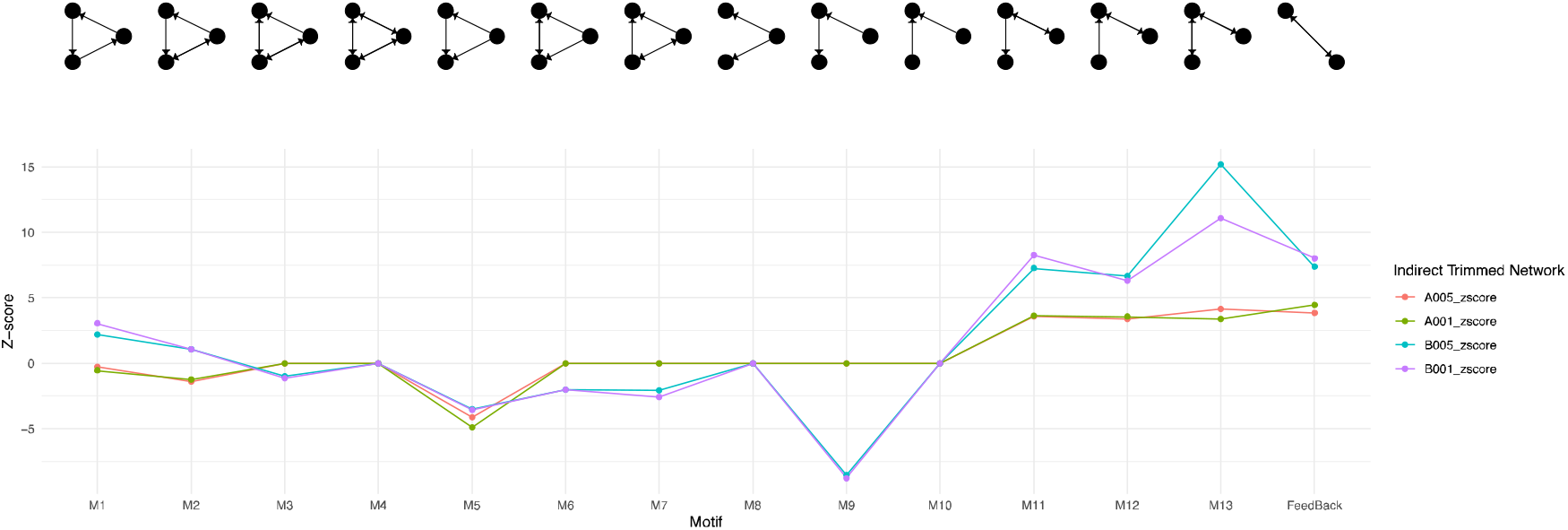
The enrichment of feedback loops in gut microbiota regulatory network. Significance of network motifs of two-node and three-node were obtained by random shuffling 10,000 times from the networks for subject A and subject B with FDR threshold of 0.01 and 0.05.

## Discussion

In this paper, we employed TENET to construct regulatory networks using longitudinal gut microbiota data from two subjects. TENET, known for its efficacy in identifying key regulatory factors in gene expression regulation from pseudotime-ordered single cell RNAseq data [26], was adapted for the analysis of time-course microbiota data. TENET was originally designed for constructing gene regulatory networks in single-cell data, its application was justified here because microbiota abundance data basically share the same data type format; read count. A challenge of this application lies in the difficulty of obtaining publicly accessible, high-quality data suitable for the application of transfer entropy. Despite of this challenge, this approach not only enabled us to measure the regulatory strength between phyla but also facilitated the identification of key regulatory genera. Notably, the top key regulatory genera were found to be associated with distinct experiences of each subject. For example, subject A exhibited *Acrobacter* as a key genus, linked to his or her experience of traveler’s diarrhea. Meanwhile, subject B had *Clostridium* identified as a key genus, associated with his or her microbiota’s persistency against *Salmonella* infection. These findings suggest the precision of TENET in capturing regulatory relationships within the microbiota.

Many studies have been discovered that *Firmicutes*, a category of intestinal microorganisms, have a substantial impact on modulating the body’s immune system [54]. During digestion, *Firmicutes* release glycoconjugates, which stimulate the production of the cytokine IL-34, leading to an enhancement of the immune response. Despite their presence throughout the body, immune balance is maintained through a feedback control mechanism. Smaller glycoconjugates are effectively managed by albumin, preventing excessive immune responses. Without these intricate mechanisms, *Firmicutes* can potentially harm vital organs. Our study provides valuable insights on which *Firmicutes* genus contribute to systemic immunity in a way that safeguards the host’s well-being. *Clostridium*, predominantly classified within the *Firmicutes* phylum, encompasses a variety of strains that are commonly found in diverse environments, including soil, sewage, the digestive tracts, and water. While some species within the *Clostridium* genus serve useful functions, others have the potential to produce toxins or cause infections, posing potential risks to the health of both humans and animals [55,56]. The original study of this time-course microbiota data suggested that Subject A’s gut microbiota reverted to its pre-travel state, whereas Subject B’s gut microbiota did not return to its pre-infection state and switched to a new stable state. Our observation of the centrality of *Clostridium* in Subject B may highlight distinct dynamics in state transitions between the two subjects.

To compare the structural difference between two subjects’ networks, we investigated three global network measures including degree distribution, modularity, and motif enrichment. We found that the degree exponent and modularity were comparable in the two networks. However, the enrichment of the two-node feedback loops and three-node motif coupled with two-node feedback loops were more enriched in the network for subject B than the network for subject A. On the other hand, cascade motif was depleted only in the network for subject B. Many studies have demonstrated that the enriched feedback loops and coupled feedback loops may be evolved for acquiring multiple steady state, hysteresis, or reliable decision making [52,57]. The differential enrichment of feedback loops between two subjects underscores their potential critical impact on varying recovery dynamics.

The examples of motif studies include not only feedback loops in biomolecular interaction networks but also trophic modules in ecological networks [35]. Specifically, motif analysis, by revealing the fundamental building blocks of ecological networks, highlights how species interactions contribute to community robustness and resilience against environmental changes [58]. These motif analyses in longitudinal microbiome studies are important, as they allow for a more understanding of microbial dynamics and the potential to predict community shifts based on observed interaction patterns.

The random rewiring of the microbiota regulatory networks of gut microbiota highlights the significant directional relationships between phyla. In both subjects, regulations within *Firmicutes* or *Proteobacteria* are markedly enriched, while regulations between these two phyla are significantly reduced. This implies a preference of microbiota to interact within genera of the same phylum rather than genera outside of that phylum.

In conclusion, this study leveraged TENET for constructing regulatory networks of longitudinal gut microbiota data from two subjects. The adapted approach allowed precise measurement of regulatory strength between phyla and identification of key regulatory genera linked to subject-specific experiences. Our findings provide insights into the roles of *Firmicutes*, especially *Clostridium*, in modulating immunity and highlight distinct dynamics in gut microbiota state transitions between subjects. These insights, drawn from data on two subjects, suggest that TENET hold promise for application in the analysis of large-scale time series microbiota datasets.

## ACKNOWLEDGMENTS

This work was supported by the National Research Foundation of Korea, funded by the Ministry of Science and ICT (NRF-2020R1A2C1005956) to JL and Basic Science Research Program through the National Research Foundation of Korea (NRF) funded by the Ministry of Education (2021R1A6A1A10044154) to JK.

