## Supplementary figures for "Inference of Causal Interaction Networks of Gut Microbiota Using Transfer Entropy"

^*^

^†^

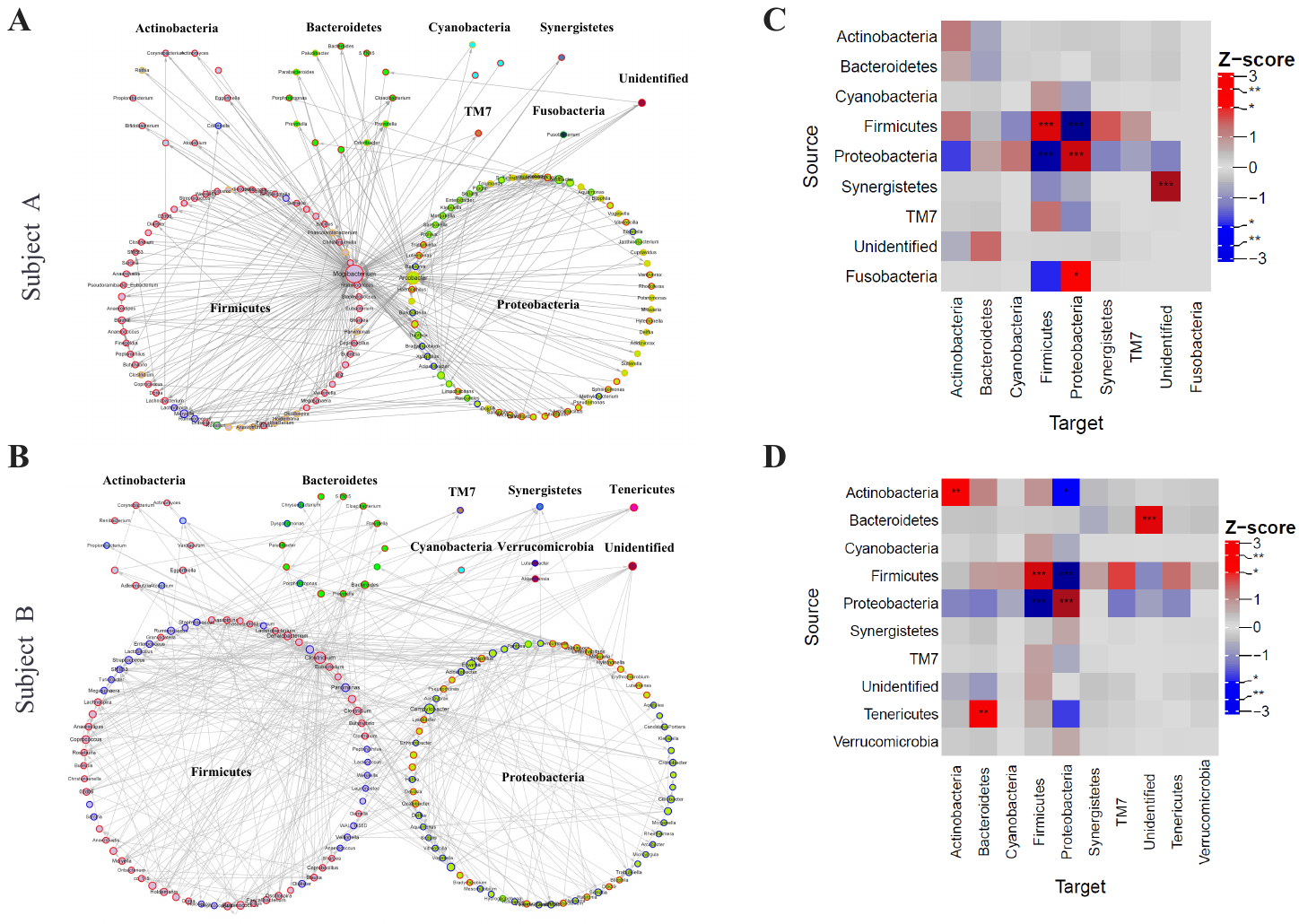

**Supplementary Figure 1. Random rewiring analysis of gut microbiota regulatory networks reveals significant regulatory relationships between phyla.** (**A-B**) Network representation of gut microbiota regulatory networks for subject A (**A**) and B (**B**) with layout of grouping by phyla. The node fill color and border color represent phylum and Leiden cluster, respectively. The node size represents the out-degree. (**C-D**) Heatmaps of the z-scores of the number of regulations between phyla for subject A (**C**) and B (**D**). The networks were obtained using FDR threshold of 0.05.

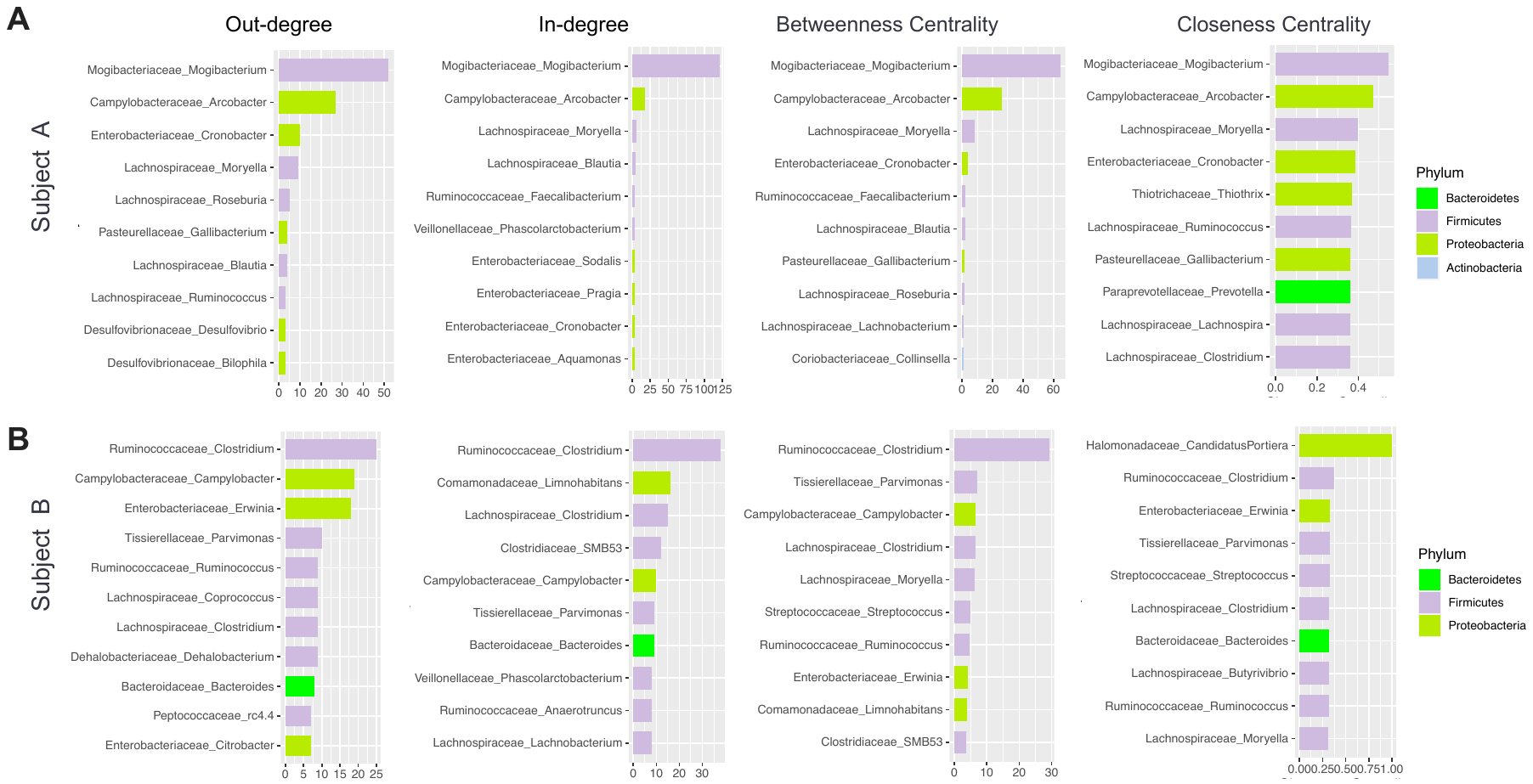

**Supplementary Figure 2. Centrality analysis of gut microbiota regulatory networks reveals candidates of key regulatory genera for regulating gut microbiota.** (**A-B**) Four centrality measures (outdegree, indegree, betweenness centrality, and closeness centrality) of subject A (**A**) and B (**B**). The color for bar represents phylum. The networks were obtained using FDR threshold of 0.05.

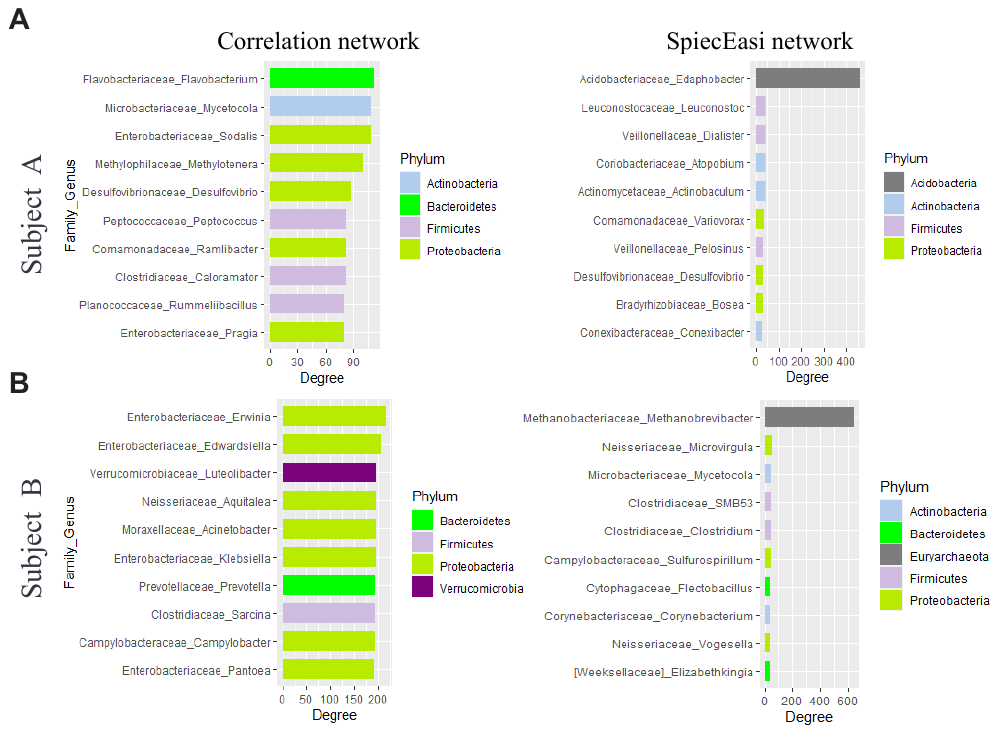

**Supplementary Figure 3. Top genus of gut microbiota regulatory networks obtained from two alternative methods.** (**A-B**) Degree of top 10 genus of correlation network and SpiecEasi network for subject A (**A**) and B (**B**). The color for bar represents phylum.

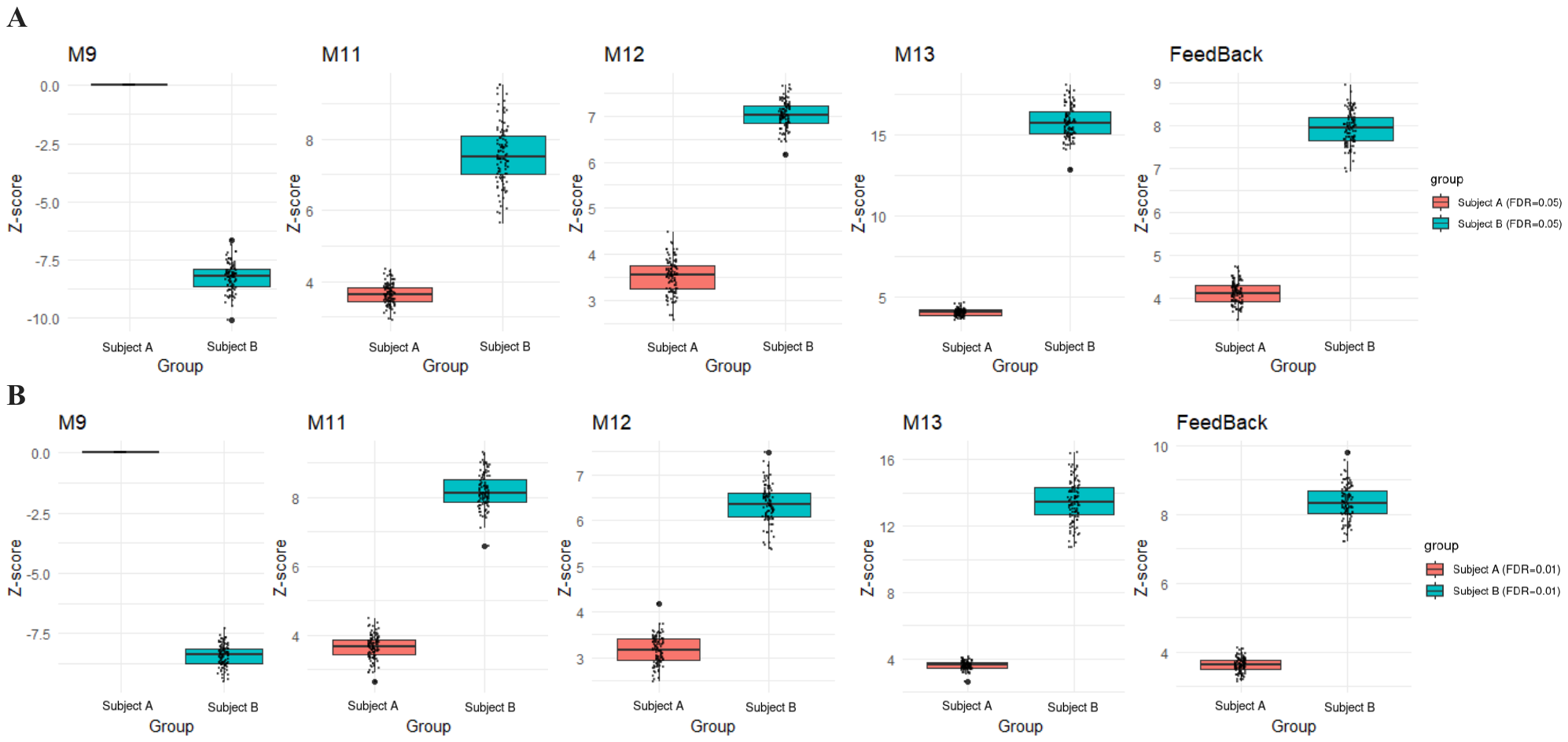

**Supplementary Figure 4. Comparison between the differential enrichment of motifs between two subjects.** (**A-B**) Z-scores of four three-node motifs including M9, M11, M12, M13, and two-node feedback when using FDR threshold of 0.05 (**A**) and 0.01 (**B**).
